# Calculating Structure Factors of Protein Solutions by Atomistic Modeling of Protein-Protein Interactions

**DOI:** 10.1101/2024.03.27.587040

**Authors:** Sanbo Qin, Huan-Xiang Zhou

## Abstract

We present a method, FMAPS(q), for calculating the structure factor, *S*(*q*), of a protein solution, by extending our **f**ast Fourier transform-based **m**odeling of **a**tomistic **p**rotein-protein interactions (FMAP) approach. The interaction energy consists of steric, nonpolar attractive, and electrostatic terms that are additive among all pairs of atoms between two protein molecules. In the present version, we invoke the free-rotation approximation, such that the structure factor is given by the Fourier transform of the protein center-center distribution function *g*_C_(*R*). At low protein concentrations, *g*_C_(*R*) can be approximated as *e*^*−βW(R)*^, where *W*(*R*) is the potential of mean force along the center-center distance *R*. We calculate *W*(*R*) using FMAPB2, a member of the FMAP class of methods that is specialized for the second virial coefficient [Qin and Zhou, J Phys Chem B 123 (2019) 8203-8215]. For higher protein concentrations, we obtain *S*(*q*) by a modified random-phase approximation, which is a perturbation around the steric-only energy function. Without adjusting any parameters, the calculated structure factors for lysozyme and bovine serum albumin at various ionic strengths, temperatures, and protein concentrations are all in reasonable agreement with those measured by small-angle X-ray or neutron scattering. This initial success motivates further developments, including removing approximations and parameterizing the interaction energy function.

## 1. Introduction

Nonspecific interactions between protein molecules play crucial roles in modulating the binding between native pairs in crowded cellular environments and mediating liquid-liquid phase separation. Yet, our knowledge of such interactions is still very limited. Small-angle X-ray and neutron scattering (SAXS and SANS) is a powerful technique for probing nonspecific interactions. In essence, the structure factor *S*(*q*) measured by SAXS and SANS is the Fourier transform of the pair distribution function, which in turn is determined by the interaction energy of the protein molecules. The structure factor at *q =* 0 and low protein concentration gives the second virial coefficient (*B*_2_), which is widely used for characterizing protein nonspecific interactions. Since the structure factor can be measured over a range of *q* and protein concentrations, it carries more information than *B*_2_ about the protein interaction energy. Structure factors have been determined for lysozyme [1-4], bovine serum albumin (BSA) [5, 6], and other proteins [7-16].

One challenge with SAXS and SANS is extracting the protein interaction energy from the structure factor. Indeed, the interpretation of the structure factor of lysozyme has generated controversy [1-3]. A common practice is to use a highly simplified interaction energy function and tune its parameters to fit the structure factor. One often-used energy function consists of steric repulsion and a two-Yukawa potential, because it produces an approximate analytical solution for the structure factor [17]. This energy function is

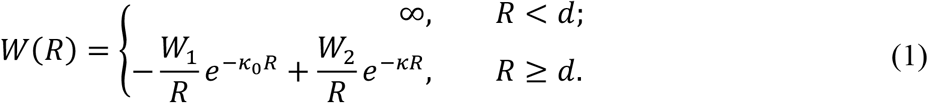

where *R* denotes the center-center distance between two protein molecules and *d* is the hard-sphere diameter. The two-Yukawa potential in the second line models short-range van der Waals attraction by the first term and long-range electrostatic repulsion by the second term. The second term can also be recognized as the Debye-Hückel potential for screened electrostatic interactions, with *κ* as the inverse of the Debye screening length. While simple models may prove adequate for fitting the structure factor, the model parameters may not provide deep physical insight into the protein under study and offer little value to other proteins.

More detailed models have been used to interpret structure factors. For example, monoclonal antibodies were modeled with a small number of fixed beads [11, 13]. Dear et al. [13] carried out molecular dynamics simulations of a 12-bead model at concentrations from 125 to 250 mg/mL to calculate the structure factors. This work marks a significant methodological advance, but because the beads were specifically designed for the monoclonal antibodies and the parameters were only tuned on these proteins, the transferability of the model and parameters to other proteins is unknown. Pasquier et al. [16] introduced a rigid coarse-grained model at the level of one bead per amino acid and carried out Monte Carlo simulations at low protein concentrations, including the exchange between two alternative conformations. Another approach is Brownian dynamics simulations of rigid proteins modeled at the all-atom level by Mereghetti et al. [18]. Other than being computationally expensive, configurational sampling by simulations and other means (see next) of atomistic proteins is the most promising approach to rigorously interpret structural factors and gain transferable physical insight into nonspecific interactions of proteins.

We have developed an approach called **f**ast Fourier transform-based **m**odeling of **a**tomistic **p**rotein-protein interactions, or FMAP [19]. FMAP was built on our previous effort in modeling protein-crowder interactions [20, 21], and calculates the interaction energy of a test protein with a protein solution [22], another molecule of the same kind as the test protein [19], or a protein molecule of a different kind [23]. These calculations yield the chemical potential for determining the binodal for liquid-liquid phase separation (FMAP*μ*) [22, 24], the second virial coefficient (FMAPB2) [19], and the cross second virial coefficient (FMAPB23) [23], respectively. In the FMAP class of methods, the sampling over the positions of the test protein is implemented through fast Fourier transform (FFT). Specifically, we express the dependence of the interaction energy on the test protein position as a sum of three-dimensional correlation functions and evaluate the correlation functions via FFT.

Here, we extend FMAPB2 into a method for predicting the structure factor. From FMAPB2 sampling of a pair of atomistic protein molecules, we obtain the potential of mean force *W*(*R*) and the protein diameter *d* determined by the steric part of the interaction energy. We introduce a modified random-phase approximation and use *W*(*R*) and *d* to calculate the structure factor. We test the *S*(*q*) predictions for lysozyme and BSA and release the FMAPS(q) program.

## 2. Theory and Calculation

We consider a protein solution comprising *N* identical, rigid protein molecules in a volume *V*. The solvent is modeled implicitly, and hence the protein solution is practically the same as a molecular liquid. We use indices *I* and *J* to refer to the individual protein molecules and index *i* or *j* to refer to atoms in a given protein molecule. The center of protein molecule *I* is specified by the position vector **R**_*I*_; the orientation is denoted as **Ω**_*I*_, which may be specified by the three Euler angles. The position of atom *i* in molecule *I* is denoted as **r**_*Ii*_, which can be written as

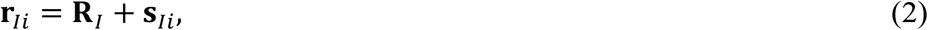

where **s**_*Ii*_ is the displacement vector from the center of molecule *I* to atom *i* in the same molecule. Typically a vector will be denoted by a bold letter, with the magnitude denoted by the same letter in italic form. Averages are calculated in the canonical ensemble.

In the presentation below, we start with definitions and relations of pair distribution functions, and then relate the scattering structure factor to pair distribution functions. Lastly we express pair distribution functions in terms of the interaction (i.e., potential) energy, and describe how to calculate the structure factor for a protein solution modeled with an atomistic interaction energy.

### 2.1. Atomic fluids

We first look at the case where the molecule is reduced to a single atom. The atom is specified solely by the position vector **R**_*I*_; **Ω**_*I*_ does not apply. The pair distribution function, *g*(*R*), is given by [25]

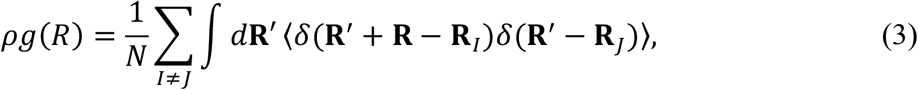

where *ρ = N/V* is the number density, *δ*(**R**) is a three-dimensional delta function, and ⟨… ⟩ denotes an ensemble average over atomic configurations. *ρg*(*R*) represents the local density of atoms around a tagged atom. The local density deviates from *ρ* due to interatomic interactions. These deviations disappear at locations far from the tagged atom; hence, at *R =* ∞, *g*(*R*) *=* 1. It is useful to introduce an alternative pair correlation function with this constant subtracted:

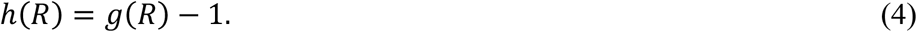

*h*(*R*) can be separated into a direct correlation function, *c*(*R*), which has a range similar to the pair potential, and indirect correlation. This separation is formally given by the Ornstein-Zernike relation

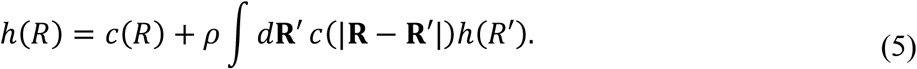

This equation can be solved only recursively for *h*(*R*). However, in Fourier space, we find

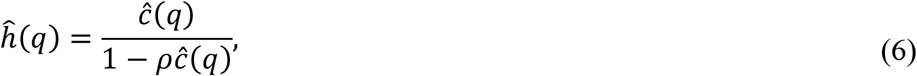

where we have used a caret “^” to denote the Fourier transform of a function.

The scattering intensity of an X-ray or neutron beam passing through the atomic fluid is given by the structure factor

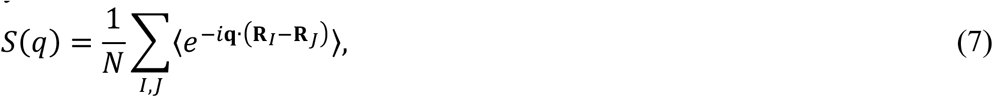

where **q** is the scattering vector. To relate *S*(*q*) to *g*(*R*), we make some manipulations:

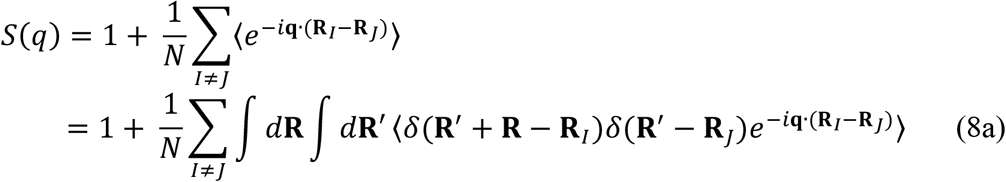

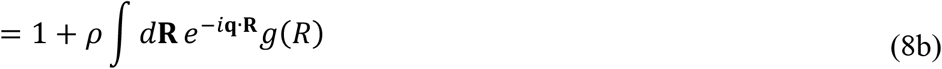

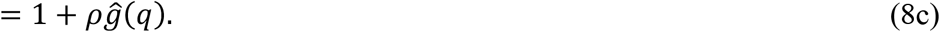

In the last two lines, the structure factor is related to the pair distribution function *g*(*R*). In terms of *h*(*R*), we get

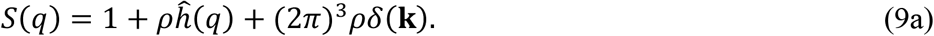

It is customary to ignore the delta-function term, which arises from scattering in the forward direction (i.e., *k =* 0 corresponds to a scattering angle of 0). Then

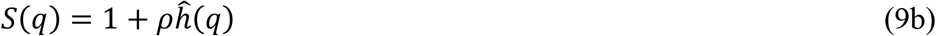

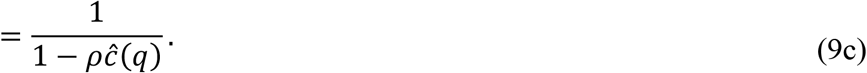

In the second step, we have used Eq. (6).

### 2.2. Molecular fluids

Analogous to the atomic pair distribution function given by Eq. (3), we can define the molecular pair distribution function [25]

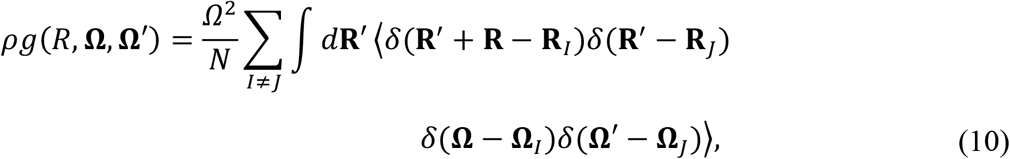

when the pair of molecules are separated by a distance *R* between the centers and adopt orientations **Ω** and **Ω**′. Here we have used the symbol *Ω* to denote ∫ *d***Ω** = 8*π*^2^. By averaging over orientations, we obtain the center-center distribution function

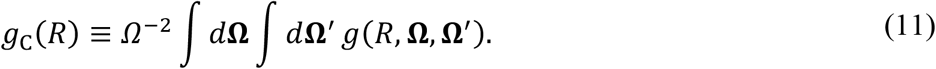

The scattering intensity due to a molecular fluid is given by [26]

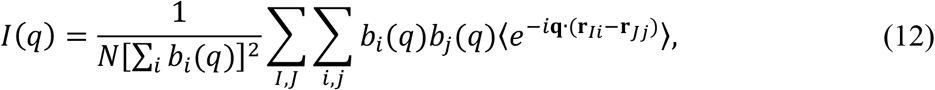

where *b*_*i*_(*q*) is the scattering length of atom *i. I*(*q*) can be separated into an intramolecular (*J = I*) term and an intermolecular (*J* ≠ *I*) term:

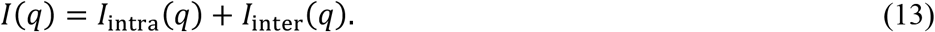

The intramolecular component is

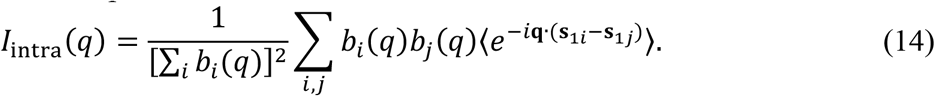

The intermolecular component is

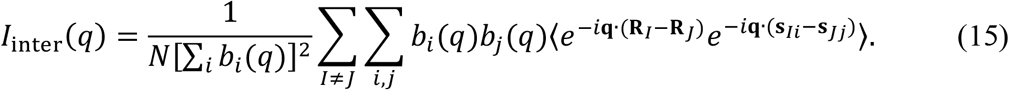

Similar to what is done in Eq. (8a), we insert delta functions to connect with the molecular pair distribution function:

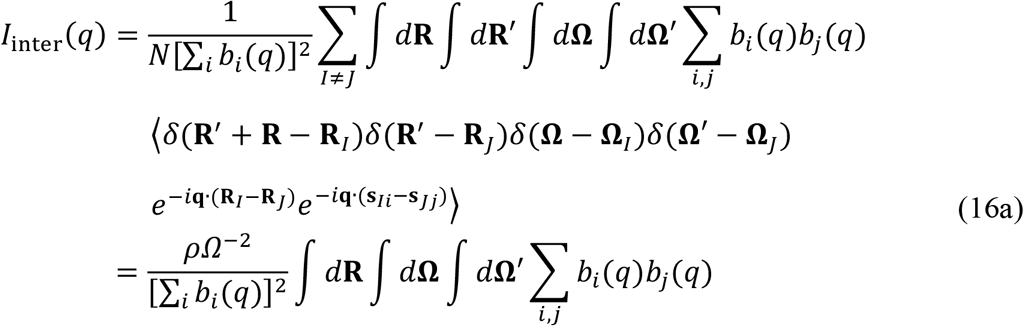

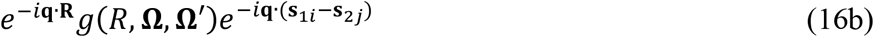

Let us first consider the *q* → 0 limit. From Eq. (14), it is obvious that *I*_intra_(0) *=* 1. Similarly, Eq. (16b) reduces to

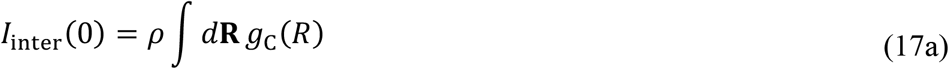

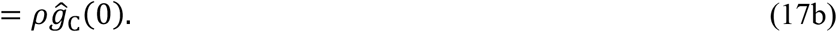

The total scattering intensity is thus

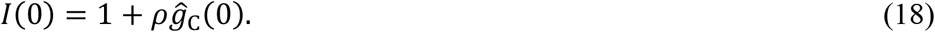

We now make approximations. The first is to orientationally pre-average *g*(*R*, **Ω, Ω**′), i.e., replace it by *g*_C_(*R*):

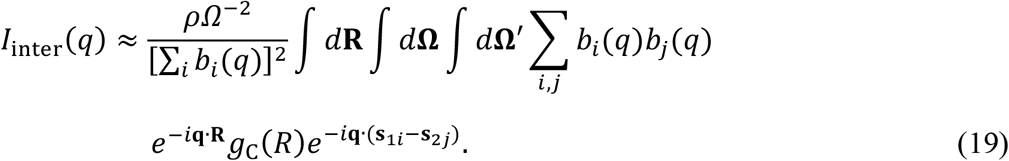

This approximation, known as free-rotation, is valid when *g*(*R*, **Ω, Ω**′) only weakly depends on molecular orientations. Next, we assume that the atomic positions and *b*,(*q*) have spherical symmetry. Due to the spherical symmetry, the ensemble average ⟨… ⟩ in Eq. (14) is unnecessary:

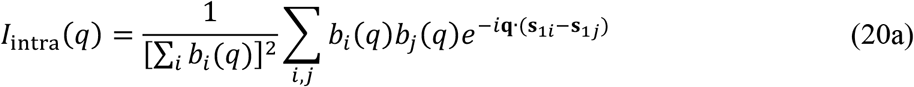

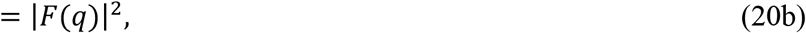

where the form factor *F*(*q*) is

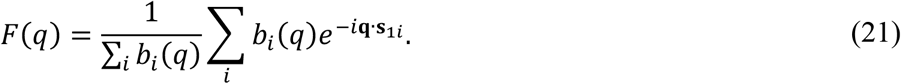

Similarly, due to spherical symmetry, for Eq. (19), the average over **Ω** and **Ω**′ is redundant. Therefore,

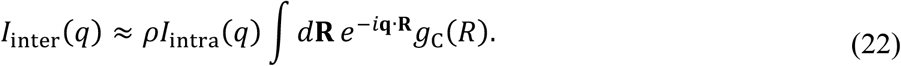

The total scattering intensity can thus be written as

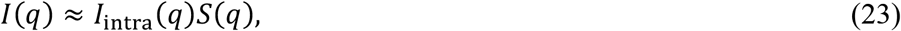

where the intermolecular structure factor is

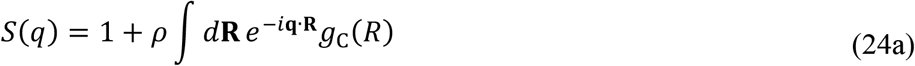

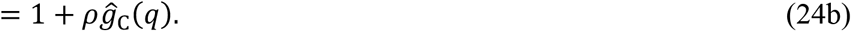

Compared with Eq. (18), we see that Eqs. (23) - (24b) are exact at the *q* → 0 limit. Again, we ignore a term containing the delta function *δ*(**k**), and find the structure factor as

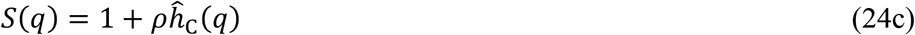

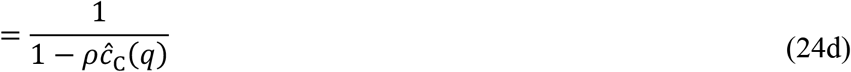

in terms of the center-center direct correlation function *ĉ*_C_(*q*).

### 2.3. Structure Factor at Low Protein Concentrations

At low molecular concentrations, the pair distribution function can be approximated by the Boltzmann factor of the pair interaction energy *U*(*R*, **Ω, Ω**′):

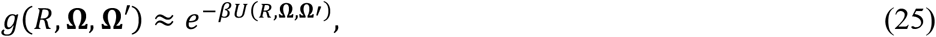

where *β =* 1*/k*_B_*T*, with *k*_B_ denoting the Boltzmann constant and *T* denoting the absolute temperature. In this subsection, the approximately equal sign “≈” specifically denotes the low-concentration limit (LCL). Correspondingly, the center-center distribution function is given by [see Eq. (11)]

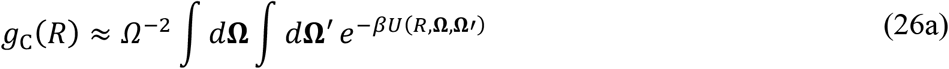

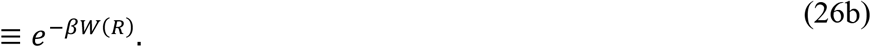

The second step serves as the definition of the potential of mean force *W*(*R*). The alternative center-center distribution function is then

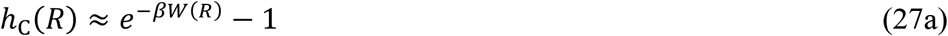

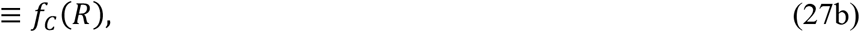

which is a Mayer function. Lastly the structure factor is [Eq. (24c)]

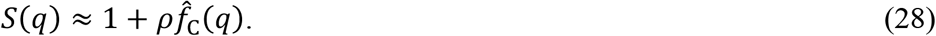

The potential of mean force can then be directly obtained from an inverse Fourier transform of the structure factor at low concentrations:

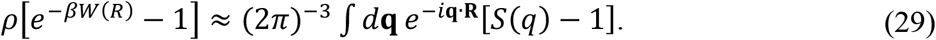

In practice, *S*(*q*) is only measured over a small range of *q* and hence the inverse Fourier transform cannot be completed.

The second virial coefficient is given by an integral of the Mayer function [19]. With the presentation notations, we have Specifically,

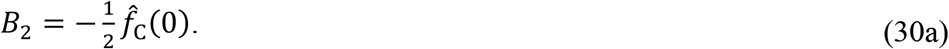

Using Eq. (28), we find

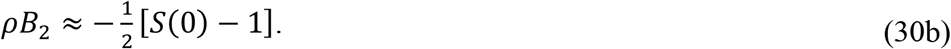

That is, *B*_2_ is equivalent to the low-concentration structure factor at a single *q* value, i.e., *q =* 0. The structure factor, even when measured over a finite *q* range, still contains more information about the potential of mean force than *B*_2_. Its dependence on molecular concentration, presented next, should provide further information about the pair interaction energy.

### 2.4. Dependence of the Structure Factor on Protein Concentration

Beyond the low-concentration limit, the pair correlation function can be calculated only numerically or by an approximation. One well-known solution is for the hard-sphere liquid under the Percus-Yevick approximation [27-29]. The pair interaction potential of hard spheres is

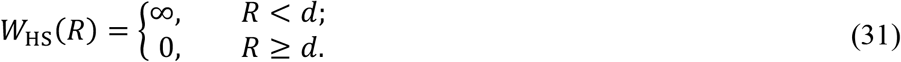

The corresponding Mayer function is

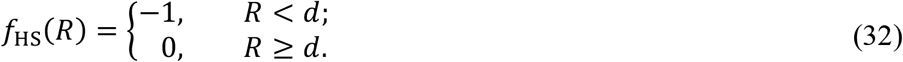

Its Fourier transform is given by

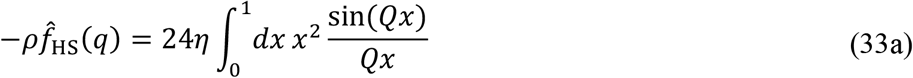

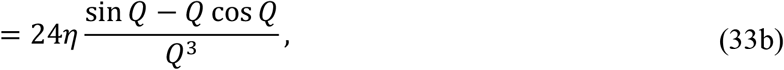

where *x = R/d, Q = qd*, and *η = πd*^3^*ρ/*6 is the volume fraction of the hard-spheres. The second virial coefficient is [Eq. (30a)]

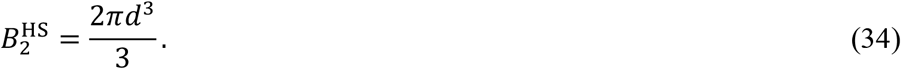

The direct correlation function *c*_HS_(*R*), according to the Percus-Yevick approximation, is

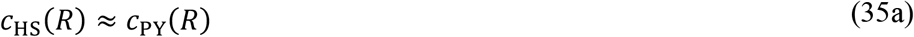

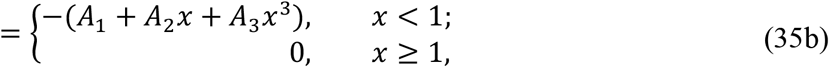

where

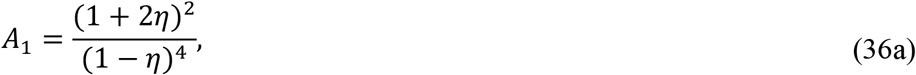

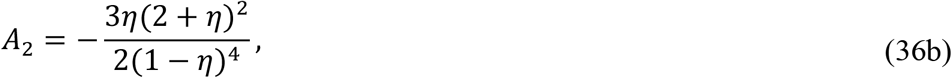

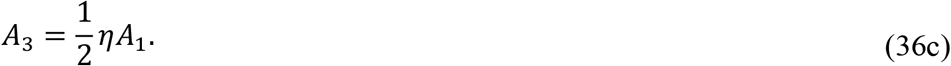

Its Fourier transform is given by

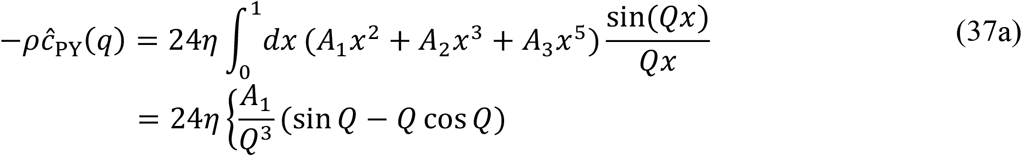

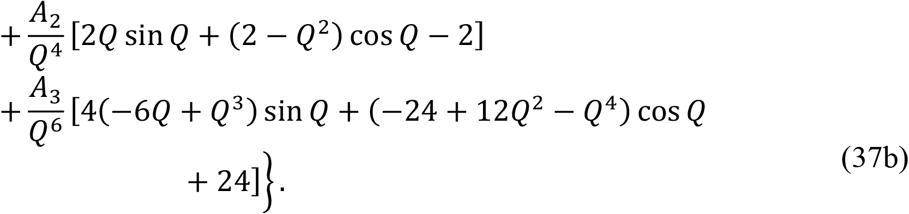

The corresponding structure factor is [Eq. (24d)]

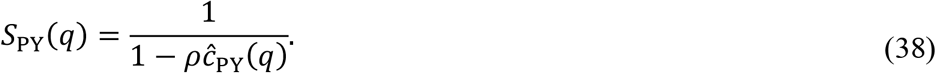

In the low-concentration limit, *A*_1_ *=* 1, *A*_2_ *= A*_3_ *=* 0; hence

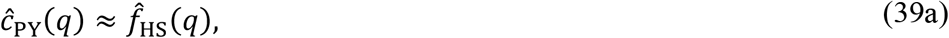

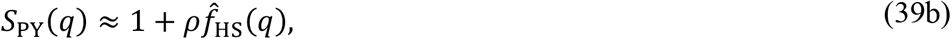

which are the correct results for *ĉ*_HS_(*q*) and *S*_HS_(*q*) in the low-concentration limit.

For a *W*(*R*) that deviates from *W*_HS_(*R*), one can obtain *c*_C_(*R*) by perturbations with the hard-sphere liquid as reference. The random-phase approximation (RPA) produces the following perturbative expression for *c*_C_(*R*) [25, 30, 31]:

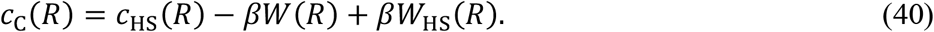

One problem with the RPA is that it does not yield the correct low-concentration limit, Eq. (28), for the structure factor. To fix this problem, we propose a modified RPA (mRPA):

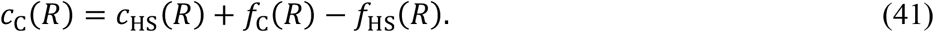

In effect, we replace *−βW*(*R*) and –*βW*_HS_(*R*) in the original RPA with the respective Mayer functions. Machin and Woodhead-Galloway [32] have noted that Eq. (41) is the correct alternative to Eq. (40) at low concentrations. Also, the Mayer function reduces to *−βW*(*R*) in the high-temperature limit (i.e., *β* → 0). Using the mRPA, Eq. (41), in Eq. (24d), we obtain

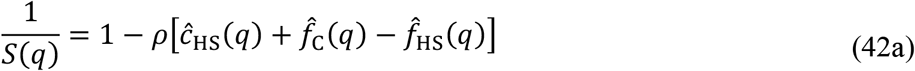

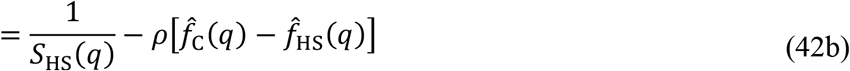

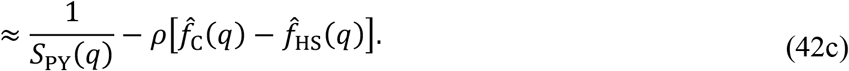

Using the low-concentration result of *ĉ*_HS_(*q*) [see Eq. (39a)] in Eq. (42a), we can easily verify that the preceding mRPA structure factor reduces to the correct result, Eq. (28), in the low-concentration limit.

Let us examine the mRPA at *q =* 0. *S*(0) is related to the isothermal compressibility

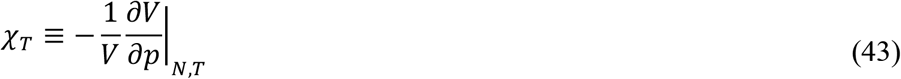

via [25]:

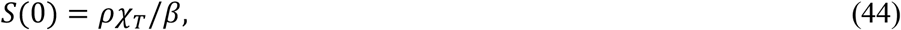

where *p* denotes pressure. Combining these two equations, we can write

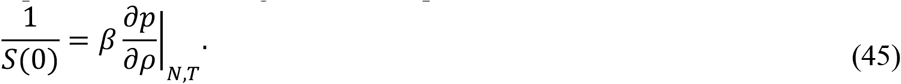

Using Eqs. (45) and (30a) in the mRPA given by Eq. (42b), we find

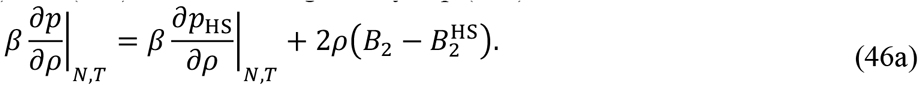

Integrating over *ρ* leads to

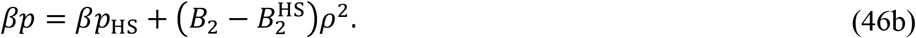

Expressing *p*_HS_ in a virial expansion, we obtain

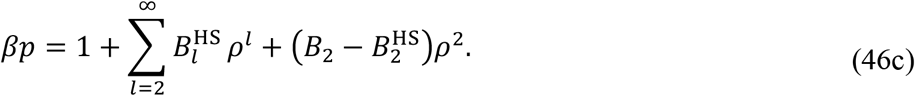

This result can be recognized as a perturbed virial expansion truncated at the second order in the number density. When the hard-sphere model is chosen to be the steric part of the real fluid, then 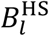 is the limiting value of the counterpart *B*_*l*_ at high temperature. As recognized by Haar and Shenker [33], starting from *l* = 3, *B*_*l*_ become close to 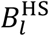 even at relatively low temperatures, therefore justifying the truncation at the second order. Our previous FMAP calculations for systems modeled with atomistic interaction functions have indeed demonstrated that the second-order term dominates over higher-order terms [34]. In short, our mRPA at *q =* 0 is equivalent to the perturbed virial expansion truncated at the second order in the number density.

### 2.5. Atomistic Interaction Energy Function and Configurational Sampling via FMAP

As in our previous studies [19, 22-24], the interaction energy, *U*(*R*, **Ω, Ω**-′), between a pair of protein molecules comprises three terms:

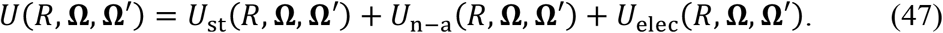

Each term is a sum over all pairs of atoms between the two molecules. The steric term is

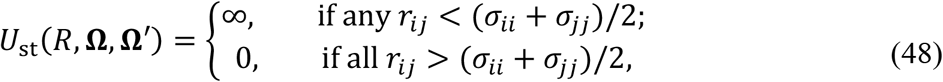

where *r*_*ij*_ denotes the distance between atom *i* in the first molecule and atom *j* in the second molecule, and *σ*_*ii*_*/*2 denotes the hard-core radius of atom *i*. The other two terms kick in only when the two molecules are clash-free [second condition in Eq. (48)]. Of these, the nonpolar attraction term, including van der Waals and hydrophobic contributions, has the form of a Lennard-Jones potential,

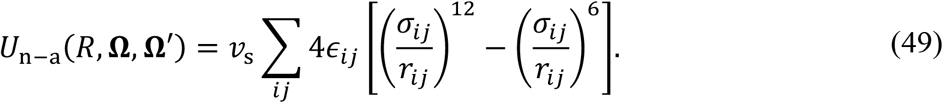

As usual, *ϵ*_*ij*_ denotes the magnitude of the attraction of the *i − j* atom pair, and *σ*_*ij*_ is the distance at which the nonpolar interaction energy is zero. The scaling constant *v*_s_ depends on the molecular mass (*M*, in kDa) of the protein molecule [19]:

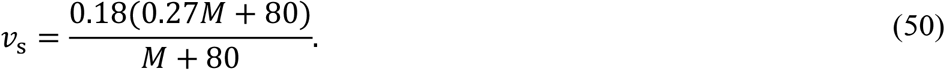

Lastly, the electrostatic term has the form of a Debye-Hückel potential,

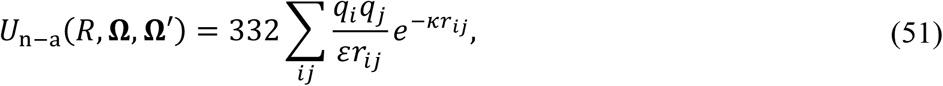

where *q*_*i*_ is the partial charge on atom *i, ε* is the dielectric constant of the solution (assumed to be water and hence decrease with increasing temperature), *κ* is the inverse of the Debye screening length and given by 2.24(*I/k*_B_*Tε*)^1*/*2^ (in Å^−1^) with *k*_B_*T* in units of kcal/mol and *I* the ionic strength in units of M. We impose a large cutoff *r*_cut_ for *r*_*ij*_ in calculating the pair interaction energy. In the previous release of the interaction energy function, *r*_cut_ was set to 36 Å [19]. Here we further increase *r*_cut_ to three times the Debye screening length when that value exceeds 36 Å (for *I* < 0.064 M at 25 ºC). There is no other parameter change in the interaction energy function.

To obtain the potential of mean force *W*(*R*), we need to sample the relative configuration of a pair of protein molecules. In Eq. (26a), this sampling is done by keeping the center-center distance at *R* and averaging over the orientations of the two molecules. In FMAP, we carry out a different but equivalent averaging procedure. We keep the center of molecule 1 at the origin and also fix its orientation, and then sample the center, **R**, of molecule 2 on a cubic grid and sample the orientation, **Ω**′, of molecule 2 uniformly by using successive orthogonal images [35]. The side length of the cubic grid is set at 2(*s*_max_ + *σ*_max_ + *r*_cut_), where *s*_max_ is the largest separation of any atom from the protein center and *σ*_max_ is the maximum of any *σ*_*ij*_, and the grid spacing is 0.6 Å in each dimension. For sampling *Ω*′, we use 4392 rotations (15º between successive rotations). From here on, the pair interaction potential will be denoted as *U*(**R, Ω**′).

We carry out the sampling in **R** by going to the Fourier space, which is the essence of FMAP and not to be confused with the Fourier transform for calculating the structure factor. In FMAP, we express the separate terms of the pair interaction energy, i.e., the steric term, the repulsive part (*r*^*−*12^) and the attractive part (*r*^*−*6^) of the nonpolar attraction term, and the electrostatic term, as correlation functions in **R**, and evaluate the correlation functions in Fourier space using FFT. An inverse FFT then produces the values of the pair interaction energy *U*(**R, Ω**′) for R at all the grid points and **Ω**′ at a particular rotation of molecule 2. We then repeat the FMAP calculations for all the 4392 rotations. To enable the expression of the *r*^*−*12^ part and the *r*^*−*6^ part as a correlation function, we have to use the geometric mean, (*σ*_*ii*_ *σ*_*jj*_)^1*/*2^, not the usual algebraic mean, as the combination rule for the *σ*_*ij*_ parameters; the geometric-mean combination rule is also used for the *ϵ*_*ij*_ parameters, as is the common practice.

### 2.6. Implementation of mRPA

Once the pair interaction energy *U*(**R, Ω**′) is obtained from the FMAP calculations, we collect the Boltzmann factor *e*^*−βU*(**R, Ω**′)^ into bins along *R*. The average of the Boltzmann factors in a bin centered at *R* gives *e*^*−βW(R)*^. Here w e use even bin intervals of 0.6 Å.

To specify the hard-sphere reference, we need to choose the diameter, *d*, of the hard spheres. To this end, we also carry out FMAP calculations for only the steric term of the interaction potential. We will use a subscript (or superscript) “st” to denote the various properties of the steric-only system. We select *d* by requiring that the hard-sphere liquid has the same second virial coefficient as the steric-only system:

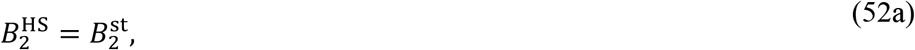

Or, equivalently,

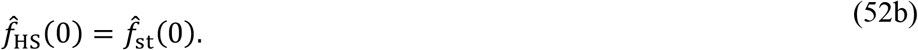

Finally, after carrying out a Fourier transform of *e*^*−βW(R)*^ *−* 1 to obtain 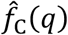, we insert it in Eq (42c) to predict the structure factor *S*(*q*).

## 3. Results and Discussion

Here we present results for the potentials of mean force for atomistic proteins, the components that go into the mRPA, and mRPA predictions. We compare these mRPA predictions with experimental data for the structural factors of two proteins, hen egg white lysozyme and BSA. We use Protein Data Bank (PDB) entries 1AKI for lysozyme [36] and 3V03 for BSA [37]. The structures of the two proteins are shown in Fig. 1. The protonation states of ionizable groups are determined from *p*Ka values predicted by PROPKA (version 3.0) [38]; a group is protonated (unprotonated) if the pH is lower (higher) than the predicted *p*Ka. The PDB coordinate file is converted into a PQR file using the PDB2PQR program (version 2.1.1) [39]; the PQR file contains atomic coordinates and partial charges. The PQR file, along with other necessary input (including the Lennard-Jones parameters *ϵ*_*ii*_ and *σ*_*ii*_, temperature, and ionic strength), is fed to the FMAPB2 code [19] to obtain the Boltzmann factor, *e*^*−βW(R)*^, of the potential of mean force *W*(*R*) and the protein diameter *d* determined [Eqs. (34) and (52a)] by the second virial coefficient, 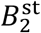, of the steric-only system. Lastly, *e*^*−βW(R)*^ and *d* are used in a new code, FMAPS(q), that calculates the structure factor by implementing the mRPA.

**Fig. 1.**
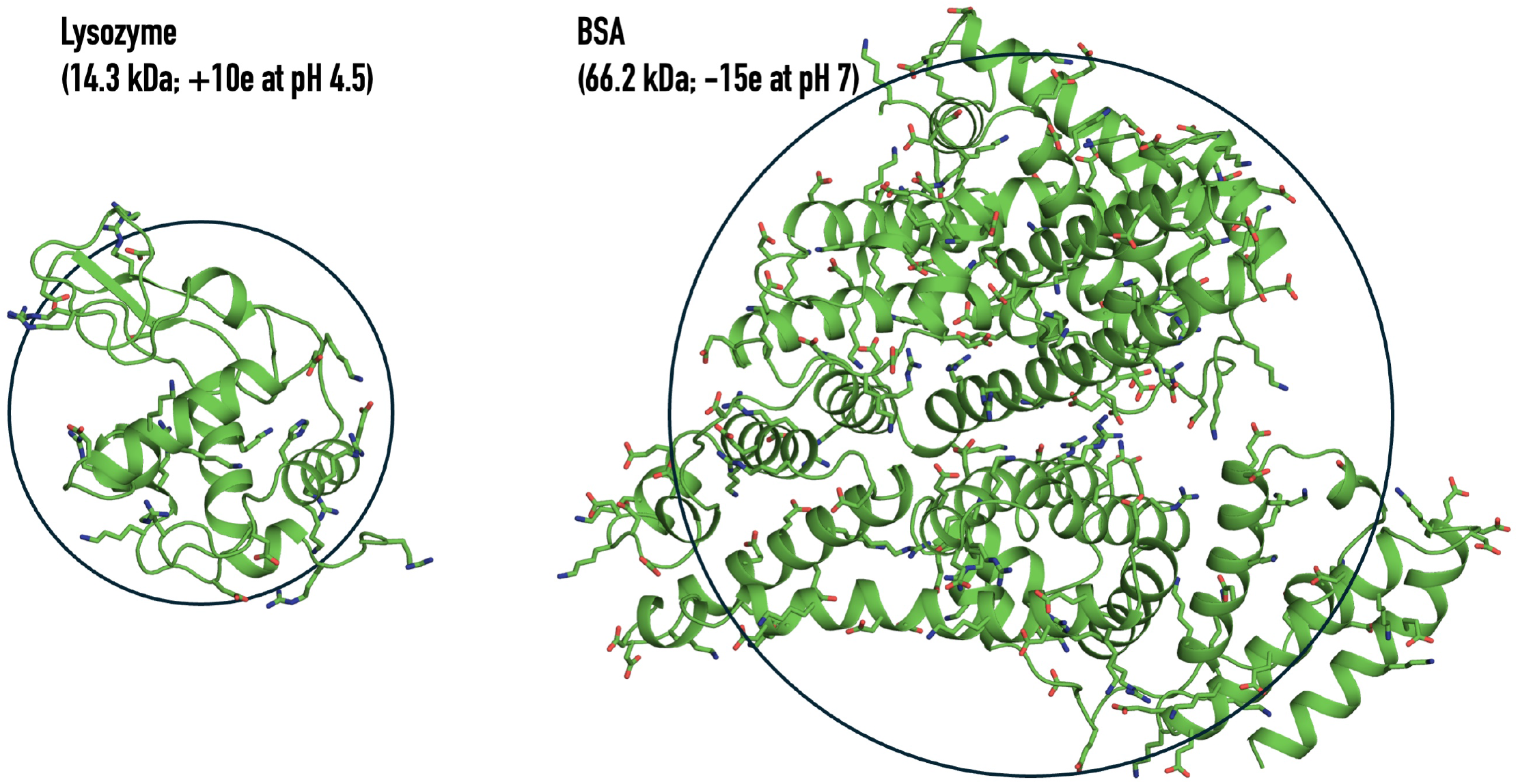
Structures of the two proteins studied here. Structures are from PDB entries 1AKI and 3V03; charged sidechains are displayed as sticks. Circles are drawn with the diameters determined by the steric part of the atomistic interaction energy function.

### 3.1. Potentials of mean force, Mayer functions, and LCL and mRPA predictions

We start with some illustrative results for lysozyme under the following solution conditions: 25 ºC temperature, pH 6.8 (net charge +8*e*), and 0.02 M ionic strength. The potential of mean force is presented in Fig. 2a in the form of a Boltzmann factor. The Boltzmann factor is less than 1 everywhere, meaning that the pair of protein molecules are repulsive to each other at all distances, due to the high net charge (+8*e*) and the low ionic strength (0.02 M). The Boltzmann factor is strictly 0 at *R* < 25 Å due to steric clash, but persists to ∼90 Å due to electrostatic repulsion.

**Fig. 2.**
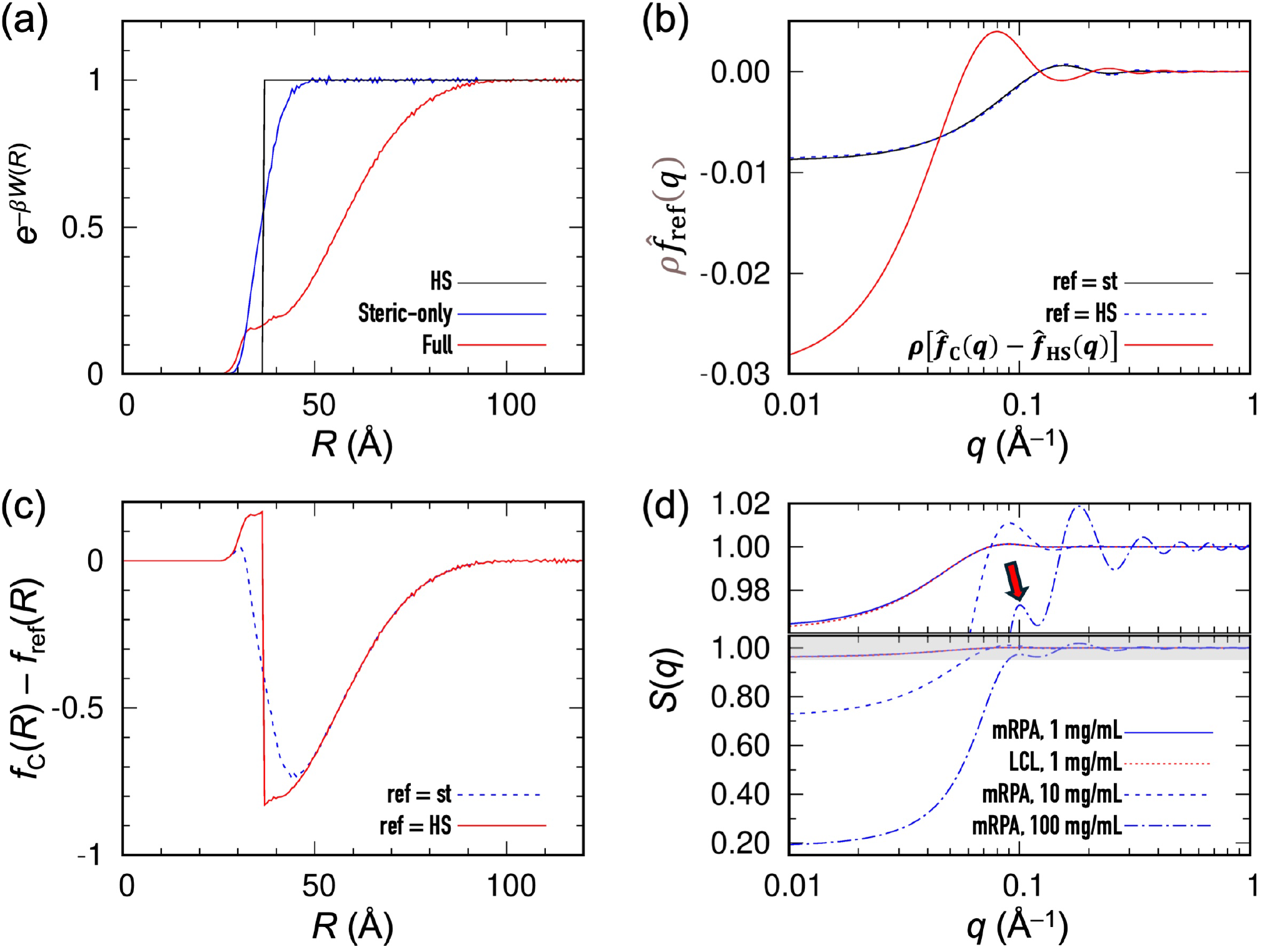
Pair potential of mean force, low-concentration limit of the structure factor, and mRPA prediction. Results are for lysozyme at 25 ºC, pH 6.8 (charge +8*e*), and *I* = 0.02 M. (a) Potentials of mean force presented in the form of Boltzmann factors, for the full interaction energy, the steric-only system, or the hard-sphere reference. (b) Fourier transforms of the Mayer functions for the steric-only system and the hard-sphere reference, or the difference in Mayer function between the full system and the hard-sphere reference. All the results are multiplied by a concentration of 1 mg/mL. (c) Differences in Mayer functions between the full system and the hard-sphere reference or the steric-only system. (d) mRPA predictions at 1, 10, and 100 mg/mL lysozyme. The prediction at 1 mg/mL is essentially identical to the low-concentration limit (LCL). The shaded top portion is also enlarged in a separate panel on top, where a red arrow indicates a low-*q* peak.

We also present the potential of mean force for the steric-only system (Fig. 2a). The corresponding Boltzmann factor switches from 0 (infinite repulsion) to 1 (no interaction) over the range from ∼30 to ∼45 Å. The gradual transition is due to our atomistic model for the protein. The protein diameter *d*, set by the second virial coefficient of the steric-only system, is 36.8 Å, roughly at the midpoint of the transition of the steric-only system. The Boltzmann factor of the corresponding hard-sphere liquid makes an abrupt transition at *R = d*. Although the Boltzmann factors of the steric-only and hard-sphere systems exhibit an obvious difference around *R = d*, their Fourier transforms, 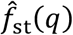 and 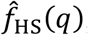, are extremely close to each other (Fig. 2b) for the small *q* values (up to 0.5 Å^−1^) accessible in typical SAXS or SANS experiments. Recall that by design, 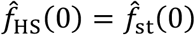 [Eq. (52b)].

What goes into the mRPA is the difference *f*_C_(*R*) *− f*_HS_(*R*) [or rather its Fourier transform; Eq. (42c)]. In Fig. 2c, we display this difference, which shows a small positive peak right before *R = d*, due to nonpolar attraction. At *R* > *d, f*_C_(*R*) *− f*_HS_(*R*) shows a large, slowly decaying negative tail due to electrostatic repulsion. In comparison, *f*_C_(*R*) *− f*_st_(*R*) shows an even smaller positive peak. We display the difference, 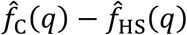, in Fig. 2b. It exhibits the first peak at *q* = 0.08 Å^−1^. This peak will become an important issue below, and hence we have ascertained its origin. It arises from the slowly decaying negative tail in *f*_C_(*R*) *− f*_HS_(*R*) (Fig. 2b). When the small positive peak of *f*_C_(*R*) *− f*_HS_(*R*) at *R* < *d* is flattened, the first peak in 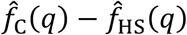 remains intact.

Lastly we display the mRPA predictions for the structure factors at lysozyme concentrations of 1, 10, and 100 mg/mL (Fig. 2d). At the lowest concentration, the mRPA result is essentially identical to the low-concentration limit, as to be expected. At both 10 and 100 mg/mL, *S*(*q*) exhibits a small peak around *q* = 0.1 Å^−1^ (indicated by a red arrow). This *S*(*q*) peak can be traced to the first peak of 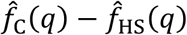, which as just stated arises from the long-range electrostatic repulsion. As the concentration increases from 10 and 100 mg/mL, the first peak of *S*(*q*) reduces in height and also moves toward higher *q*.

Below we compare mRPA predictions against experimental structure factors for lysozyme and BSA. The solution conditions for which the comparison is made are listed in Table 1.

**Table 1.** List of protein solutions.

| Temperature<br>(°C) | pH | Charge<br>(*e*) | Ionic strength<br>(M) | Protein concentration<br>(mg/mL) | Expt method<br>[refs] |
| --- | --- | --- | --- | --- | --- |
| Lysozyme (*d* = 36.8 Å) |  |  |  |  |  |
| 10 | 8 | 7 | 0.01 | 20 | SAXS [4] |
| 10 | 8 | 7 | 0.06 | 20 |  |
| 10 | 7.4 | 8 | 0.15 | 20 |  |
| 25 | 8 | 7 | 0.01 | 20 |  |
| 25 | 8 | 7 | 0.06 | 20 |  |
| 23 | 7.4 | 8 | 0.15 | 20 |  |
| 40 | 8 | 7 | 0.01 | 20 |  |
| 40 | 8 | 7 | 0.06 | 20 |  |
| 37 | 7.4 | 8 | 0.15 | 20 |  |
| 25 | 6.8 | 8 | 0.02 | 50 | SANS [3] |
| 25 | 5.3 | 10 | 0.02 | 100 |  |
| 25 | 4.5 | 10 | 0.02 | 175 |  |
| 25 | 4.5 | 10 | 0.02 | 225 |  |
| BSA (*d* = 69.4 Å) |  |  |  |  |  |
| 20 | 7 | -15 | 0.005 – 0.015 | 0.9, 4.5, 9, 18, 20, 40,<br>45, 60, 80 90, 100,<br>200 | SAXS [5, 6] |
| 20 | 7 | -15 | 0.05 | 100 | SAXS [5] |
| 20 | 7 | -15 | 0.1 | 100 |  |
| 20 | 7 | -15 | 0.3 | 100 |  |
| 20 | 7 | -15 | 0.5 | 100 |  |

### 3.2. Lysozyme: effects of ionic strength and temperature

In Fig. 3a, we display *e*^*−βW(R)*^ for lysozyme at *I* = 0.01, 0.06, and 0.15 M, with temperatures at 25 ºC, 25 ºC, and 23 ºC, respectively, and pH (charge) at 8 (+7*e*), 8 (+7*e*), and 7.4 (+8*e*), respectively, to match with the SAXS experimental conditions of Tanouye et al. [4]. Compared to the result in Fig. 2a at *I* = 0.02 M, the repulsive tail at *I* = 0.01 M extends to an even longer distance, ∼120 Å, due to the increased Debye screening length. As the ionic strength is raised to 0.06 M, the electrostatic repulsion is screened enough such that *e*^*−βW(R)*^ at *R* = 32.1 Å rises above 1, corresponding to an attractive *W*(*R*); the potential of mean force is still repulsive at longer distances, but both the amplitude and the range are considerably curtailed. At *I* = 0.15 M, the peak of *e*^*−βW(R)*^ at *R* = 32.1 Å climbs even further as the nonpolar attraction term becomes dominant there, and the repulsive tail all but disappears.

**Fig. 3.**
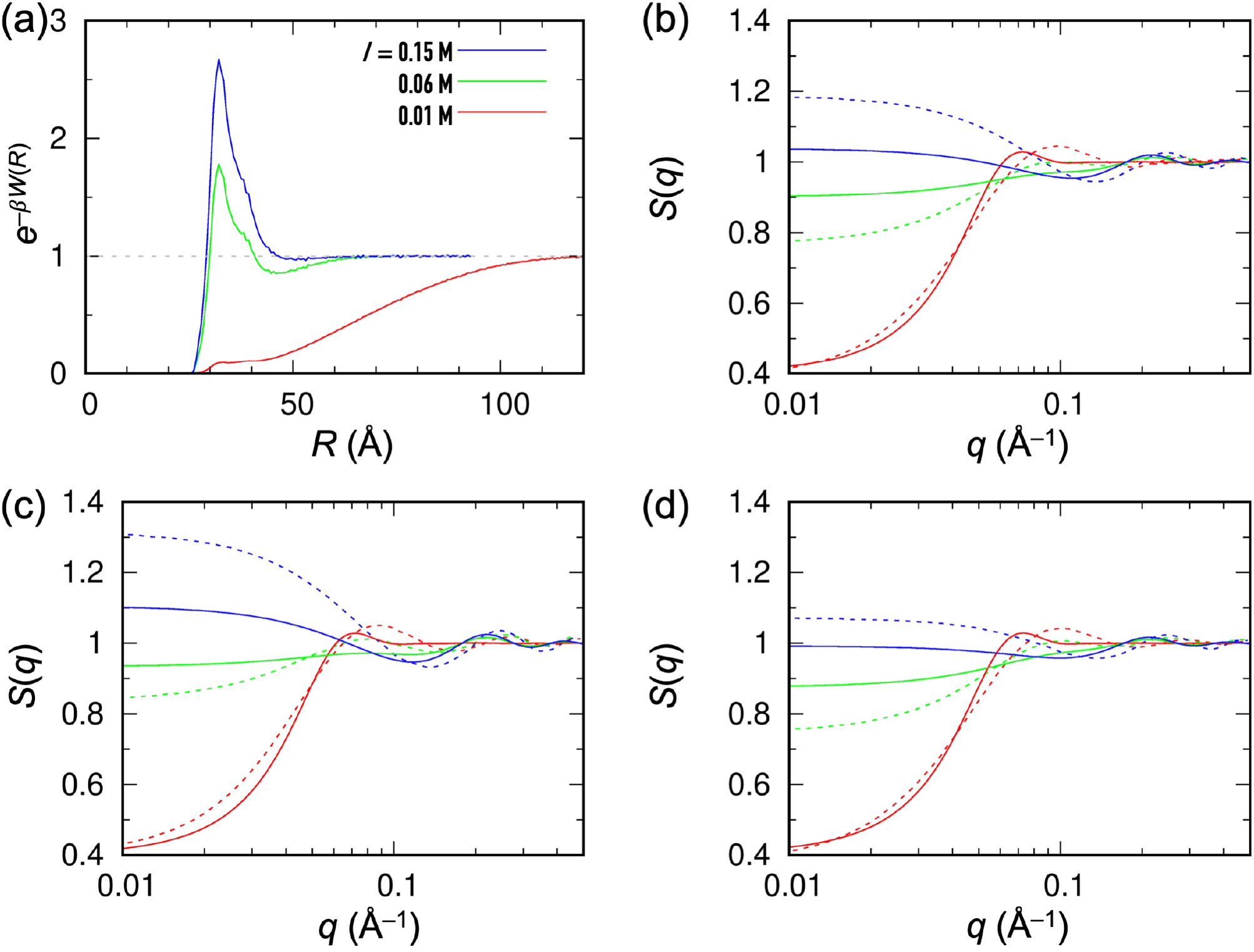
Effects of ionic strength and temperature on the structure factor of lysozyme. (a) Boltzmann factors at *I* = 0.01, 0.06, and 0.15 M. The temperatures are at 25, 25, and 23 ºC, respectively; the pH values (net charges) are 8 (+7*e*), 8 (+7*e*), and 7.4 (+8*e*), respectively. (b) Comparison of mRPA (solid curves) and experimental (dashed curves) structure factors; the latter are from SAXS measurements by Tanouye et al. [4]. (b) Corresponding results at 10 ºC. (c) Corresponding results at higher temperatures (40, 40, and 37 ºC).

We compare in Fig. 3b the predicted and measured structural factors for lysozyme at 20 mg/mL under the above conditions. The results match well at *I* = 0.01 M, both showing a *S*(*q*) value at *q* = 0.01 Å^−1^ that is substantially below 1, which we can attribute to the strong electrostatic repulsion between a pair of protein molecules. At *I* = 0.06 M, both the measured and predicted structural factors move toward a uniform value of 1, which we can attribute to the cancelation of contributions from the attractive part of the potential of mean force around *R* = 32.1 Å and the repulsive part at longer distances. The mRPA prediction does underestimate the deviation of *S*(*q*) from 1 at *q* = 0.01 Å^−1^, indicating that the net repulsive effect from our potential of mean force is too weak. At *I* = 0.15 M, both the measured and predicted *S*(*q*) at *q* = 0.01 Å^−1^ rise above 1, which we can attribute to the dominance of the nonpolar attraction term in the pair interaction potential. However, the effect of the nonpolar attraction is underestimated.

Next we present the effects of temperature. The corresponding plots at 10 ºC are displayed in Fig. 3c and those at 40 ºC for *I* = 0.01 and 0.06 M and at 37 ºC for *I* = 0.15 M are displayed in Fig. 3d. The experimental results for *S*(*q*) at *q* = 0.01 Å^−1^ exhibit two trends.

First, *S*(*q*) increases with decreasing temperature. Second, the temperature effects are minuscule at *I* = 0.01 M, modest at *I* = 0.06 M, and prominent at *I* = 0.15 M. The mRPA predictions recapitulate both of these trends. Note that the structure factor of the hard-sphere reference has no temperature dependence [see Eq. (37b)]. In our model, temperature affects the solvent dielectric constant *ε* and also appears in the Debye screening length and directly in the Boltzmann factors. For electrostatic interactions, *T* and *ε* appear as a part of the product *k*_B_*Tε* both in the Debye screening length and in the Boltzmann factor. Because *ε* decreases with increasing temperature, to a crude estimation *k*_B_*Tε* remains a constant and therefore temperature does not change the electrostatic contribution. At *I* = 0.01 M where electrostatic repulsion dominates, we thus expect little temperature effects, as is the case in both our mRPA prediction and the experimental data. At *I* = 0.15 M where nonpolar attraction dominates, temperature affects the potential of mean force only through its direct appearance in the Boltzmann factor. In this case, the Boltzmann factor of the potential of mean force and therefore the structure factor increase with decreasing temperature, just as seen in our mRPA prediction and the experimental data.

### 3.3. Lysozyme over a concentration range

The results in Fig. 3b-d are for a single lysozyme concentration of 20 mg/mL. Now we present mRPA results for lysozyme over a range of concentrations and compare them with the SANS data of Liu et al. [3]. The comparison is displayed in Fig. 4a-d for 50, 100, 175, and 225 mg/mL lysozyme, respectively. The solution conditions are: 25 ºC temperature, 0.02 M ionic strength, and pH 6.8 (charge +8*e*) for 50 mg/mL lysozyme and pH 5.3 to 4.5 (charge +10*e*) for the three higher lysozyme concentrations.

**Fig. 4.**
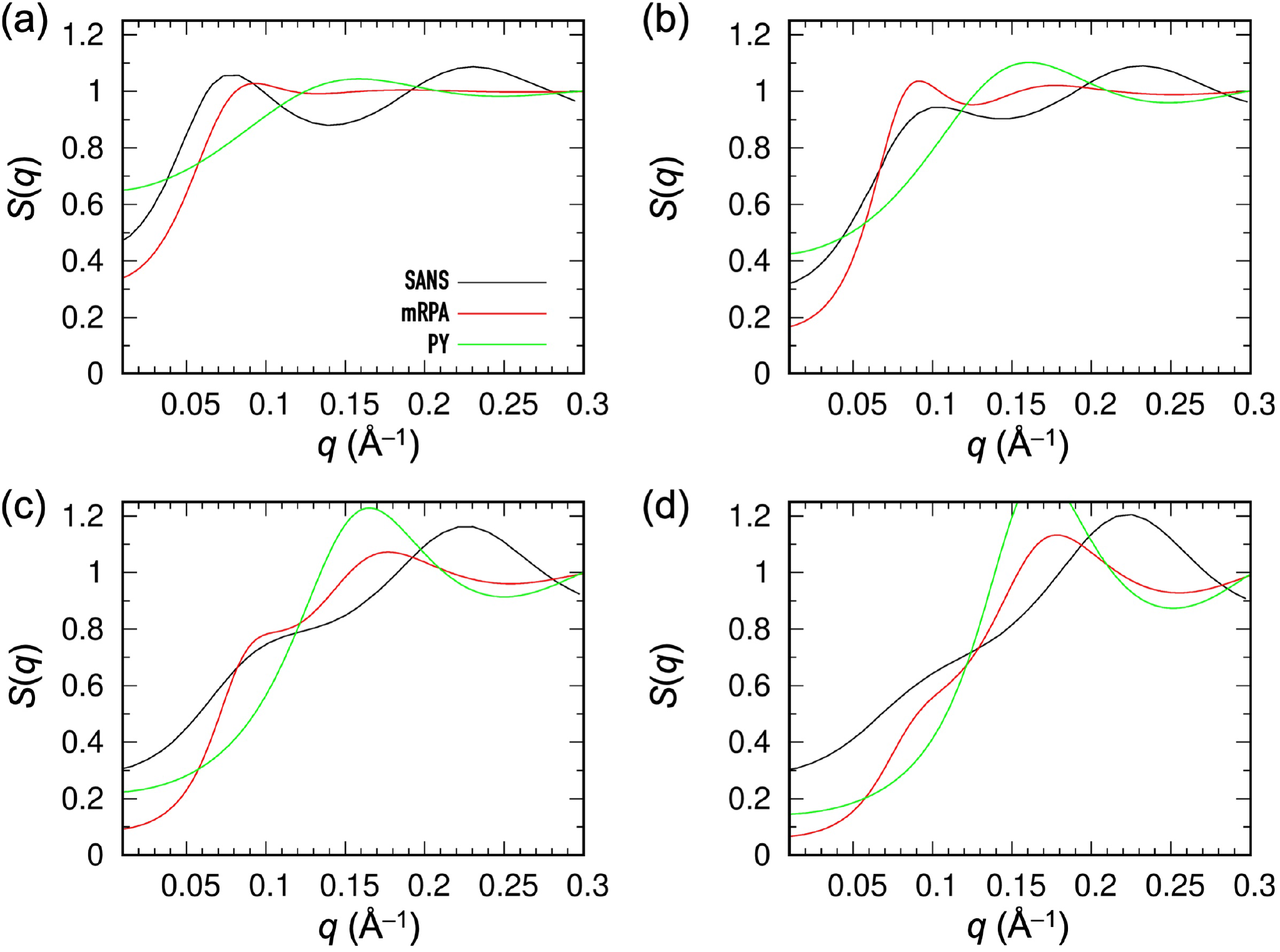
Dependence of lysozyme structure factors on concentration. (a) 50 mg/mL. (b) 100 mg/ml. (c) 175 mg/mL. (d) 225 mg/mL. The temperature is 25 ºC, ionic strength is 0.02 M, and pH values (net charges) are 6.8 (+8*e*), 5.3 (+10*e*), 4.5 (+10*e*), and 4.5 (+10*e*). The mRPA predictions, Percus-Yevick structure factors, and SANS measurements [3] are shown as red, green, and black curves, respectively.

The most prominent feature in the experimental *S*(*q*) is a first peak near *q* = 0.1 Å^−1^ at 50 and 100 mg/mL lysozyme that turns into a shoulder at the two higher lysozyme concentrations. As alluded to in Introduction, this low-*q* peak has generated much controversy. Stradner et al. [1] claimed that this peak is due to the formation of molecular clusters. Both Shukla et al. [2] and Liu et al. [3] pushed back, showing that potentials of mean force comprising a short-range attractive term and a long-range repulsive term could produce a low-*q* peak. Shukla et al. specifically attributed this peak to the repulsive term, whereas Liu et al. related the low-*q* peak to the second peak in the center-center pair distribution function (around *R =* 2*d*). In complete agreement with the experimental *S*(*q*), the mRPA predictions exhibit the low-*q* peak at 50 and 100 mg/mL lysozyme and a shoulder at the two higher lysozyme concentrations. As explained above when presenting Fig. 2d, the low-*q* peak arises solely from the long-range electrostatic repulsion at the low ionic strength of 0.02 M.

The experimental *S*(*q*) data also exhibit a second peak at *q* = 0.23 Å^−1^, with the peak height increasing at higher protein concentrations. To identify the origin of this second peak, we include in Fig. 4 the Percus-Yevick structure factor for the hard-sphere reference system, which shows a single peak at *q* = 0.17 Å^−1^ (corresponding to 2*π/d*) with increasing height. This peak in *S*_GH_(*q*) is related to the contact peak of the center-center pair distribution function, and results in the second peak of the mRPA structure factor. The latter peak position is at a *q* value that is too low compared to the experimental counterpart. It could be moved to the correct position by a small decrease in the hard-sphere diameter *d*, from 36.8 Å to 30 Å. The need for this decrease may be due to the nonspherical shape of lysozyme, which allows two protein molecules to approach each other at distances shorter than *d* (see Fig. 2a), or perhaps reflects our treatment of the protein molecules as rigid. Molecular flexibility would reduce the distance of closest approach.

### 3.4. BSA: effects of ionic strength

In Fig. 5a, we display *e*^*−βW(R)*^ for BSA at ionic strengths ranging from 0.005 to 0.5 M, with temperature at 20 ºC and pH at 7 (charge = –15*e*), to match with the SAXS experimental conditions of Zhang et al. [5]. At *I* = 0.005 M, the Boltzmann factor is indicative of strong repulsion, with a tail extending to ∼200 Å. As the ionic strength is increased, the electrostatic repulsion becomes more and more screened, and the Boltzmann factor rises more and more rapidly, with the tail retracting further and further toward smaller *R*. At *I* = 0.3 M, the Boltzmann factor rises above 1, as the nonpolar attraction term begins to dominate at short range, and the repulsive tail disappears. At *I* = 0.5 M, the attractive peak further grows in amplitude. The BSA diameter determined by the steric part of the interaction potential is 69.4 Å.

**Fig. 5.**
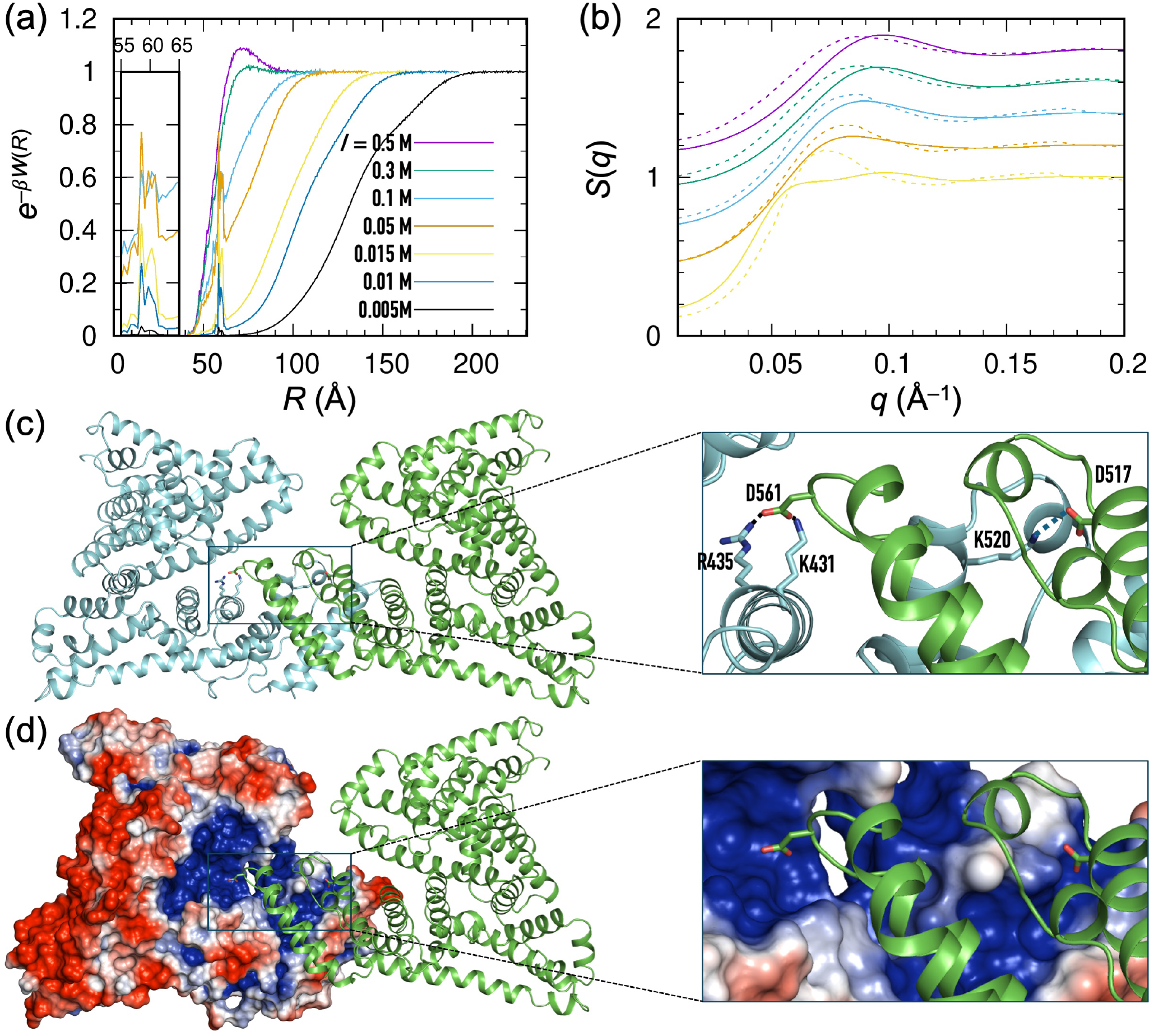
Effects of ionic strength on BSA pair interactions and structure factor. (a) Boltzmann factors at *I* = 0.005, 0.01, 0.015, 0.05, 0.1, 0.3, 0.5 M. The local peak at *R* = 60 Å is also shown in a zoomed view. Temperature = 20 ºC; pH = 7 (charge –15*e*). (b) Comparison of mRPA (solid curves) and SAXS (dashed curves) [5] structure factors at a concentration of 100 mg/mL and ionic strengths from 0.015 to 0.5 M [in colors matching those in (a)]. Curves are shifted in the vertical direction to avoid overlap. (c) A preferential complex from the FMAPB2 sampling, stabilized by local electrostatic and nonpolar attraction. A zoomed view shows salt bridges between two BSA molecules. (d) The electrostatic potential of BSA shows a positive region in a mainly negatively charged surface. A zoomed view shows that the positive patch is lined by negatively charged residues of the second BSA molecule.

In Fig. 5b we display the mRPA structure factors for 100 mg/mL BSA in the ionic strength range of 0.015 to 0.5 M. The predictions match well with the SAXS data of Zhang et al. There is some mild underestimation of the structure factors at the highest ionic strengths, perhaps hinting at the need for strengthening the nonpolar attraction. Comparing Fig. 3b and Fig. 5b, we see that whereas for lysozyme an ionic strength of 0.15 M is able to turn the *S*(*q*) value at *q* = 0.01 Å^−1^ from less than 1 to greater than 1, for BSA the *S*(*q*) value at *q* = 0.01 Å^− 1^ remains at less than 1 even at *I* = 0.5 M. The mRPA calculations reveal two reasons for this difference. Firstly, the net charge of BSA (–15*e*) has a much greater magnitude than the counterpart for lysozyme (+8*e*); therefore electrostatic screening requires a much higher ionic strength for BSA. Moreover, the SAXS measurement for BSA was made at a much higher concentration (100 mg/mL) than the counterpart for lysozyme (20 mg/mL); that means that steric repulsion plays a much greater role in the case of BSA, which helps keep *S*(0) below 1.

The Boltzmann factors also exhibit a sharp peak at *R* = 60 Å for ionic strengths up to 0.1 M. This peak corresponds to preferential complexes formed between two BSA molecules (Fig. 5c). These complexes are stabilized by both local electrostatic and nonpolar attraction, afforded by charge complementarity and shape complementarity, respectively (Fig. 5d). While these preferential complexes may not make a significant contribution to the mRPA structure factor of BSA presented here, their roles as competitors to protein-protein native complexes [40] and as the drive for liquid-liquid phase separation [41] have been revealed. Characterizing these roles requires atomistic modeling.

### 3.5. BSA over a concentration range

Both Zhang et al. [5] and Heinen et al. [6] reported structure factors of BSA at different concentrations. These SAXS measurements were made with the protein in water (without buffer). In their modeling by Brownian dynamics simulations, Mereghetti et al. [18] assumed an ionic strength of 0.005 M to account for the dissolution of CO2, residual salt, and the dissociation of surface groups from the protein [6]. Here we further assume that the ionic strength increases to 0.01 M at protein concentrations of 40 and 45 mg/mL and further to 0.015 M at higher protein concentrations. The latter value has already appeared as the lowest ionic strength in Fig. 5a. In Fig. 6a, we compare our mRPA structure factors, those predicted by Mereghetti et al., and the SAXS data of Heinen et al. for BSA at concentrations from 0.9 to 90 mg/mL. The predictions of Mereghetti et al. are pretty good, but ours are in even closer agreement with the SAXS data.

**Fig. 6.**
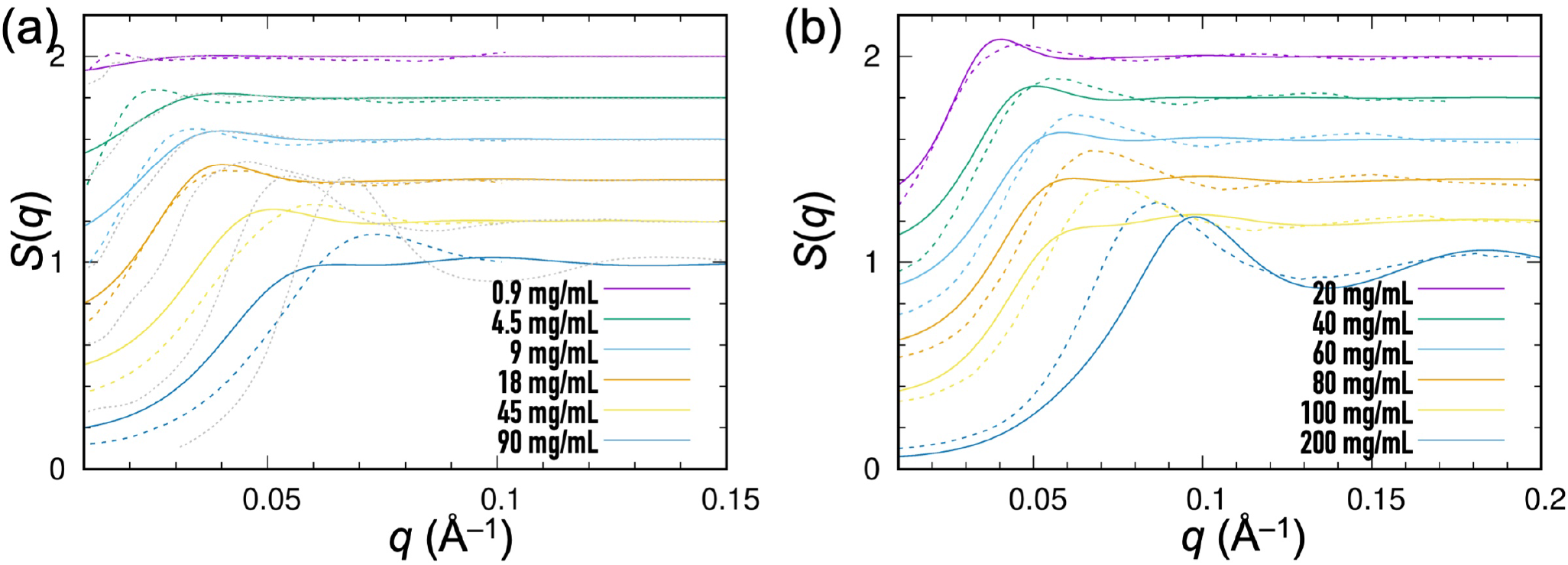
Dependence of BSA structure factors on concentration. (a) Comparison of mRPA (solid curves) and Brownian dynamics [18] (dotted curves) predictions and SAXS structure factors (dashed curves) [6] at concentrations from 0.9 to 90 mg/mL. (b) Comparison of mRPA (solid curves) and SAXS structure factors (dashed curves) [5] at concentrations from 20 to 200 mg/mL. Curves are shifted in the vertical direction to avoid overlap.

Good agreement is also achieved between our mRPA predictions and the data set of Zhang et al., which covers the concentration range from 20 to 200 mg/mL. These authors also reported the structure factor at 400 mg/mL. However, such a high concentration corresponds to a nominal volume fraction of 0.64, at which the Percus-Yevick approximation breaks down for the hard-sphere reference system and hence the mRPA cannot be implemented.

### 3.6. Software release

We have implemented FMAPS(q) into a web server, accessible at https://zhougroup-uic.github.io/FMAPSq/. Users input the potential of mean force in the form of the Boltzmann factor *e*^*−βW(R)*^ and the protein diameter *d*, both of which are produced by the FMAPB2 program, and obtain the structure factor at the desired protein concentration.

At the web server page, users can also download the JavaScript code for FMAPs(q), **fmapsq.js**, and the FMAPB2 package. After clicking the “registration to download software” button, users are directed to a link, where they can register and download. We recommend that users run the FMAPs(q) code through the cross-platform JavaScript runtime environment, nodejs (https://nodejs.org). The command is

$ **nodejs** [*dir/*]**fmapsq.js** *Bltz*.*txt diam qstrt qstep Nq Cmol*

The meanings of the parameters are: *dir*, the directory where **fmapsq.js** is saved; *Bltz*.*txt*, a text file containing *e*^*−βW(R)*^ as a function of *R*; *diam*, the protein diameter; *qstrt*, the starting *q* value for outputting *S*(*q*); *qstep*, the step size for incrementing *q*; *Nq*, the number of *q* values; and *Cmol*, the protein concentration in mM. Alternatively, the protein concentration can be input in units of mg/mL. In that case, the parameter “*Cmol*” in the command line is replaced by “*Cmas MW*”, where *Cmas* represents the mass concentration in mg/mL, and *MW* is the molecular mass in kDa.

The core of our FMAPB2 package was written in C, calling FFTW, OpenMP, and Zlib libraries. It also requires python2, bash, awk, and optionally gnuplot and ghostscript for graphics. The FMAPB2 package can directly run in a Linux environment that satisfies these requirements, by the command

$ [*dir/*]**fmapb2** *pro*.*pqr ionc temp nthrd*

The meanings of the parameters are: *dir*, the directory where the FMAPB2 files are saved; *pro*.*pqr*, the PQR file for the protein; *ionc*, the ionic strength in M; *temp*, the temperature in ºC; and *nthrd*, value for OMP_NUM_THREADS in OPENMP, representing the number of threads (*≤* maximum number of hardware threads) to use for parallel regions of the FMAPB2 C code.

To remove the dependence on the specific Linux environment, users can take advantage of Apptainer (formerly Singularity; https://github.com/apptainer/apptainer), a container platform designed for mobility of compute [42, 43]. With Apptainer, the FMAPB2 package can run on any modern Linux distribution that is installed on bare metal or a Virtual Machine in Windows or Mac. After installing Apptainer, users first generate a system image file fmapsys.sif, by the command

$ **apptainer** build --fakeroot *fmapsys*.*sif fmapsys*.*def*

The definition file fmapsys.def is provided in the FMAPB2 download. The FMAPB2 package can then be run by the command

$ **apptainer** exec *fmapsys*.*sif* [*dir/*]**fmapb2** *pro*.*pqr ionc temp nthrd*

## 4. Conclusion

We have presented the FMAPS(q) method that combines atomistic modeling with the mRPA to predict the structure factors of protein solutions. Without adjusting any parameters, the predictions match well with SAXS and SANS data for two proteins under a wide range of solution conditions and protein concentrations, demonstrating robustness. FMAPS(q) calculations unequivocally show that the low-*q* peak of the lysozyme structure factor is due to long-range repulsion. Importantly, FMAPS(q) is based on modeling at the pair level and avoids expensive simulations of concentrated protein solutions. We have implemented FMAPS(q) into a web server and released the source code, at https://zhougroup-uic.github.io/FMAPSq/.

With this initial success of FMAPS(q), we can consider further developments. Two directions are worth noting. The first is to remove some of the approximations that go into the mRPA. For example, instead of hard spheres, a better reference system for protein solutions may be hard particles with a spheroidal shape. The second direction is to tune parameters in our pair interaction energy function. In particular, both Fig. 3b and Fig. 5b indicate possible improved agreement with experimental data if the scaling constant *v*_Q_ is increased somewhat. A systematic effort in parameterization, involving SASA/SANS data for more proteins, promises to further improve the accuracy of FMAPS(q).

Compared to simple models like the two-Yukawa potential, our atomistic model offers distinct values. First, the model is transferrable from protein to protein, i.e., it can be used for different proteins without case-by-case parameterization. Second, the atomistic model is more fundamental in nature and its parameters are more physically meaningful, and hence it produces more insightful interpretations of structural factor data. For example, FMAPS(q) is able to capture the temperature dependence of the lysozyme structure factor without introducing ad hoc temperature-dependent parameters. Lastly, our atomistic model produces preferential complexes such as that shown in Fig. 5c. SAXS/SANS may not be sensitive enough to detect these preferential complexes, but other techniques such as NMR can [40].

Still, SAXS/SANS measurements can help parameterize our atomistic model, which can then be used to characterize preferential complexes in activities ranging from inhibition of protein-protein native complexes to mediation of liquid-liquid phase separation [40, 41].

### CRediT author contribution statement

**Sanbo Qin:** Conceptualization, Data curation, Formal analysis, Investigation, Software, Visualization. **Huan-Xiang Zhou:** Conceptualization, Data curation, Investigation, Supervision, Validation, Visualization, Writing-original draft, Writing-review & editing.

## Data and code availability

Source data for making the plots reported in figures are posted on GitHub at https://github.com/hzhou43/FMAPSq. The FMAPS(q) webserver and source code as well as the FMAPB2 package can be downloaded at https://zhougroup-uic.github.io/FMAPSq/.

## Acknowledgments

This work was supported by Grant GM118091 from the National Institutes of Health.

